## Supplementary material for "Investigating the role of urban vegetation alongside other environmental variables in shaping *Aedes albopictus* presence and abundance in Montpellier, France": S1_Text

**S1 Text: Detailed description of bivariate and multivariate analysis.**

All details of the bivariate and multivariate analyses are provided in the R scripts 4_model_building and 6_multivariate_model_building, respectively. S2 Figure summarizes the entire analysis workflow. For bivariate analyses, data were centered but not scaled. A hurdle modeling approach was used: on the one hand, a GLMM with a binomial distribution and a logit link, with presence (1) or absence (0) of Aedes albopictus as the response variable, and trap location nested within the sampling area and the sampling session as random intercepts. On the other hand, a strictly abundance model was fitted with a truncated negative binomial distribution and a log link, with the number of Ae. albopictus caught per trap per day as the response variable, and the same random effects (trap location within sampling area and sampling session). Bivariate analyses were conducted using the **glmmTMB** package. Marginal R² was calculated for each model using the **performance** package.

Meteorological, landscape, microclimatic, air quality, and socio-economic variables with higher R² marginal were selected. A first selection was realized for variables with p-value > 0.20. The correlations between variables were assessed using the Pearson Correlation Coefficient > 0.7 and VIF < 3.

Then, Random Forest models were trained for presence (binary classification) and abundance (classification regression) separately, using the **ranger** method from the **caret** package. First, to reduce the number of predictors, recursive feature elimination (RFE) was performed on all subsets of 1 to 10 selected variables. The best subset was chosen based on cross-validated performance using spatial cross-validation (leave-one-area-out), with classification accuracy for presence models and mean absolute error (MAE) for abundance models. Next, presence and abundance models were optimized with a random grid search of 10 combinations and trained with 500 trees. The minimum node size was set to 1 for presence models and 5 for abundance models. The mtry values were 3 and 2, respectively. The splitting rule was set to ‘extratrees’ for presence models, which applies extremely randomized splits to increase tree diversity and reduce overfitting. For abundance models, the splitting rule was ‘variance’, which selects splits that minimize the variance of the response within nodes to improve prediction accuracy. Model performance was evaluated using AUC for presence models and MAE for abundance models. Both spatial (leave-one-area-out) and temporal (leave-one-session-out) cross-validation were applied, i.e., training on all data except one area or session and testing on the left-out fold. Cross-validated ROC results for presence models are presented in S7 Fig. For abundance models, cross-validated MAE results, stratified by Aedes albopictus abundance intervals, are presented in S8 Fig.

We assessed spatial autocorrelation of residuals from both presence and abundance Random Forest models using the **spatialRF** package at various distances (0–250 m) with Moran’s I. For the presence model, Moran’s I values ranged from -0.044 to -0.015, with corresponding p-values between 0.188 and 0.662, indicating no significant spatial autocorrelation. For the abundance model, Moran’s I values were slightly negative (-0.072 to -0.045) and p-values ranged from 0.051 to 0.278. No spatial autocorrelation was statistically detected.
