## Supplementary material for "Investigating the role of urban vegetation alongside other environmental variables in shaping *Aedes albopictus* presence and abundance in Montpellier, France": S1_Table

| **Name of Area** | **Abb.** | **Number and identification of traps** | **Surface (in m2 (Ha))** | **% of landcover** | **Flora** | **Fauna** | **Nearest sampling area and distance (trap to trap)** |
| --- | --- | --- | --- | --- | --- | --- | --- |
| Park of Aiguelongue | PRK-AGL | 2 (BG01, BG02) | 9357 (0.94) | Low veget: 18.64%  High veget: 80.89%  Others: 0.47%  Buildings: 0%  Roads : 0% | Mediterranean vegetation, mainly grass, pine trees, laurel shrubs and a few bamboos. Play areas. | Humans, domestic carnivores, squirrels, birds ( | IMP-SCU (1415 m) |
| Botanical Garden | PRK-BOT | 3 (BG03, BG04, BG05) | 53603 (5.4) | Low veget: 29.58%  High veget: 51.05%  Others: 15.18%  Buildings: 3.39%  Roads: 0.80% | A park with a bamboo grove, aromatic plants, vegetable garden, bromeliad greenhouse, water features, lawn and trees like oak, cipress and magnolia. | People, cats, frogs, goldfish, bats, various birds (ducks, heron, pigeon, turtle dove), squirrels, etc. | IMP-BBI  (115m) |
| Lemasson | RES-LEM | 2 (BG11, BG14) | 32036 (3.2) | Low veget: 28.9%  High veget: 22.5%  Others: 22.6%  Buildings: 18.5%  Roads: 7.5% | Residential gardens with lawns, fruit trees, ivy and ornamental plants in pots | Human, cats, dogs , birds | RES-AGR  (1420m) |
| Aiguerelles | RES-AGR | 2 (BG15, BG16) | 34549 (3.5) | Low veget: 26.5%  High veget: 14.2%  Others: 23.0%  Buildings: 20.7%  Roads: 15.6% |  | Human, cats, dogs , birds, chickens | RES-LEM (1420m) |
| Soulas | RES-SOUL | 2 (BG12, BG13) | 33415 (3.341) | Low veget: 34.0%  High veget: 15.3%  Others: 21.1%  Buildings: 16.9%  Roads: 12.7% |  | Human, cats, dogs , birds | IMP-ACA (2160m) |
| Saint-Charles University | IMP-SCU | 1 (BG21) | 5880  (0.580) | Low veget: 18.0%  High veget: 36.0%  Others: 45.0%  Buildings: 0 %  Roads: 1.0% | Car park with plane trees, a few aromatic plants, ivy and laurel | Humans, domestic carnivores, squirrels, birds | PRK-BOT (230m) |
| Bouisson Bertrand Institute | IMP-BBI | 1 (BG22) | 1215 (0.12) | Low veget: 3.1%  High veget: 7.8%  Others: 48.2%  Buildings: 40.9%  Roads : 0% | Yard hedge of laurel, ivy and plane trees | Human, birds | PRK-BOT (115m) |
| Diderot Institute | IMP-DID | 1 (BG23) | 1001 (0.1) | Low veget: 0%  High veget: 0%  Others: 47.10%  Buildings: 51.8%  Roads: 1.10% | Terraced with two olive trees | Human, birds | PRK-BOT (250m) |
| Hotel Acapulco | IMP-ACA | 1 (BG24) | 497 (0.05) | Low veget: 30.6%  High veget: 31.3%  Others: 11.5%  Buildings: 19.4%  Roads: 7.2% | Yard with bamboo hedges, acanthus and abelia grandiflora | Humans, domestic carnivores, squirrels, birds | PRK-BOT (290m) |
