## Supplementary material for "Investigating the role of urban vegetation alongside other environmental variables in shaping *Aedes albopictus* presence and abundance in Montpellier, France": S2_Table

| **Variable topics** | **Name** | **Abbreviation** | **Unit** | **Data Processing** | **Source of data set or acquisition method** | **Spatial/temporal scales of source data** | **Format of dataset** | **Link to download the data set** |
| --- | --- | --- | --- | --- | --- | --- | --- | --- |
| Microclimatic | Local hourly minimum, maximum, and average temperatures | tmin, tmax, tmean | Celsius degree (°C) | **Real-time and 48h-lagged variables.**  Hourly microclimatic data were recorded during the sampling period, covering the 24 hours prior to collection, as well as 48 hours before. For example, the average relative humidity during collection would be named Rhmean_collection. | Field data acquisition with data loggers (Hygro button) | Point data (traps) ; Hourly | NA | NA |
|  | Local hourly minimum, maximum, and average relative humidity | rhmin, rhmax, rhmean | % |  |  |  |  |  |
| Meteorological | Daily minimum, maximum, and average temperatures | TMIN, TMAX, TMEAN | Celsius degree (°C) | **Weeks-lagged variables.** The weekly TMIN, TMAX and TMEAN were calculated for different time lags ranging from 0 to 6. For example, the average temperature for weeks 2 to 4 collected would be named TMEAN_2_4. | ODEE daily meteorological dataset for the Hérault department (one station in Montpellier) | Point data (station) ; Hourly | CSV | <https://odee.herault.fr/index.php/thematiques/climatologie> |
|  | Growing Degree Day | GDD | Celsius degree (°C) | **Weeks-lagged variables.** The weekly cumulative GDD/CUMRF was calculated for different time lags ranging from 0 to 6. For example, the GDD for weeks 2 to 4 collected would be named GDD_2_4. |  |  |  |  |
|  | Daily Cumulated Rainfall | CUMRF | mm |  |  |  |  |  |
|  | Precipitations during sampling, 24 h and 48 h before | Precipitation | mm | **Real-time and 48h-lagged variables.**  The total rainfall during the day of sampling, 24 h and 48 h before, was recorded. |  |  |  |  |
|  | Daily average Wind Speed | WS | meter/second | **Weeks-lagged variables.** The weekly average RH/WS was calculated for different time lags ranging from 0 to 6. For example, the average wind speed for weeks 2 to 4 collected would be named WS_2_4. | Meteorological dataset from the Frejorgues airport station, belonging to the Meteo France national meteorology agency network | Point data (station) ; 3-hours | CSV | <https://meteo.data.gouv.fr/datasets/donnees-climatologiques-de-base-quotidiennes/> |
|  | Daily average Relative Humidity | RH | % |  |  |  |  |  |
|  | Daily minimum, maximum, average atmospheric pressure and difference of pressure during sampling, 24 h and 48 h before | Patmin, Patmax, Patmean, Patdiff | Pascal (Pa) | **Real-time and 48h-lagged variables.** The minimum, maximum, and mean atmospheric pressure were extracted for the sampling period and the preceding 24 and 48 hours, along with the mean daily pressure difference between these periods. For example, the average atmospheric pressure during collection would be named Patmean_collection |  |  |  |  |
| Quality air | Concentration of nitrogen monoxyde | NO | µg/m^3^ | **Weeks-lagged variables.** The weekly average concentration was calculated for different time lags ranging from 0 to 6. For example, the average concentration of O3 for weeks 2 to 4 collected would be named as O3_2_4. | Dataset of the Daily measurement of the Main pollutants in Occitanie, from six stations belonging to Atmo Occitanie, the regional air quality monitoring association | Point data (stations) ; Daily | Shapefile | <https://data-atmo-occitanie.opendata.arcgis.com/datasets/> |
|  | Concentration of nitrogen dioxide | NO2 |  |  |  |  |  |  |
|  | Concentration of Particulate matter with diameters < 10 µm | PM10 |  |  |  |  |  |  |
|  | Concentration of Particulate matter with diameters < 2.5 µm | PM2.5 |  |  |  |  |  |  |
|  | Concentration ozone | O3 |  |  |  |  |  |  |
| Land cover | Percentage of low vegetation (< 3m height) | % low veget | % | Calculated for 4 buffers (20m, 50m, 100m and 250m) based on trap locations. For example, the percentage of low vegetation at 50 m would be named % low veget 50 m. | the fine-scale vegetation dataset created in 2019 by Montpellier Mediterranée Métropole | Vector data (polygon > 1 m²); 2019 | Shapefile | <https://data.montpellier3m.fr/dataset/vegetation-fine-2019> |
|  | Total edge length of low vegetation (< 3m height) | Edge_low_vegetation | m |  |  |  |  |  |
|  | The average patch size of low vegetation (< 3m height) | Area_low_vegetation | Ha |  |  |  |  |  |
|  | Number of patches of low vegetation (< 3m height) | Patches_low_vegetation | Number of patches |  |  |  |  |  |
|  | Percentage of high vegetation (> 3m height) | % high veget | % |  |  |  |  |  |
|  | Total edge length of high vegetation (> 3m height) | Edge_high_vegetation | m |  |  |  |  |  |
|  | The average patch size of high vegetation (> 3m height) | Area_high_vegetation | Ha |  |  |  |  |  |
|  | Number of patches of high vegetation (> 3m height) | Patches_high_vegetation | Number of patches |  |  |  |  |  |
|  | Percentage of buildings | % buildings | % |  | The French national database of Buildings (BDNB) | Vecor data (polygon); 2023 | Geopackage | <https://www.data.gouv.fr/fr/datasets/base-de-donnees-nationale-des-batiments/> |
|  | Total edge length of buildings | Edge_buildings | m |  |  |  |  |  |
|  | The average patch size of buildings | Area_buildings | Ha |  |  |  |  |  |
|  | Percentage of roads | % roads | % |  | Urban Atlas of the Copernicus European Union’s space program component | Vector data (polygon>= 2500 m^2^) | Geopackage | <https://land.copernicus.eu/en/products/urban-atlas/urban-atlas-2018> |
|  | Total edge length of roads | Edge_roads | m |  |  |  |  |  |
|  | The average patch size of roads | Area_roads | Ha |  |  |  |  |  |
|  | Percentage of others | % others | % |  |  |  |  |  |
|  | Land cover diversity | Shannon diversity index (*H*) | NA | $H=- \sum_{i=1}^{S} p_{i}\ln(p_{i})$, where $p_{i}$ is the proportion of land cover type *i* within the buffer and *S* is the total number of land cover types. | Calculated from the three previous land cover databases | NA | NA | NA |
| Demographic | Human population | Pop | Number of inhabitants | Sum of population for 4 buffers (20m, 50m, 100m and 250m) based on trap locations. | The Montpellier fine-scale population dataset, created in 2016 by the Languedoc-Roussillon Geographic Information System (SIG-LR) | Vector data (points per building) ; 2020 | Shapefile | <https://data.montpellier3m.fr/dataset/distribution-fine-de-la-population> |
| Socio-economics | Number of poor households | Pov_Hous | Number of poor households | Intersection between trap locations and INSEE grids: Association of traps with the number of poor households and the percentage of buildings constructed before 1945, between 1945 and 1970, between 1970 and 1990 and after 1990. | French National Institute for Statistics and Economic Studies (INSEE) | Raster data (200m) ; 2019 | Geopackage | https://www.insee.fr/fr/statistiques/7655515 |
|  | Number of homes built before 1945 | Hom_Be_45 | Number of buildings |  |  |  |  |  |
|  | Number of homes built between 1945 and 1970 | Hom_45_70 |  |  |  |  |  |  |
|  | Number of homes built between 1970 and 1990 | Hom_70_90 |  |  |  |  |  |  |
|  | Number of homes built after 1990 | Hom_Af_90 |  |  |  |  |  |  |
