## Supplementary material for "Investigating the role of urban vegetation alongside other environmental variables in shaping *Aedes albopictus* presence and abundance in Montpellier, France": S3_Table

| **Trap** | **Landscape metrics** | **In 20 m buffer** | **In 50 m buffer** | **In 100m buffer** | **In 250 m buffer** |
| --- | --- | --- | --- | --- | --- |
| BG01 | % low vegetation | 25.95 | 30.17 | 17.38 | 19.84 |
|  | % high vegetation | 73.39 | 55.27 | 50.92 | 36.88 |
|  | % buildings | 0 | 2.80 | 9.28 | 15.20 |
|  | % roads | 0 | 4.12 | 10.30 | 9.64 |
|  | % others | 0.66 | 7.64 | 12.12 | 18.44 |
|  | Total edge length of low vegetation (m) | 85.5 | 724.5 | 2920.5 | 26266.5 |
|  | Total edge length of high vegetation (m) | 76.5 | 569 | 2410 | 16862.5 |
|  | Total edge length of buildings (m) | 0 | 74 | 854.5 | 9267 |
|  | Total edge of roads (m) | 0 | 132 | 1284.5 | 8493 |
|  | Average size of a patch of low vegetation (Ha) | 0.017 | 0.040 | 0.013 | 0.013 |
|  | Average size of a patch of high vegetation (Ha) | 0.095 | 0.219 | 0.115 | 0.078 |
|  | Average size of a patch of buildings (Ha) | 0 | 0.006 | 0.023 | 0.030 |
|  | Average size of a patch of roads (Ha) | 0 | 0.033 | 0.109 | 0.126 |
|  | Number of patches of low vegetation | 2 | 6 | 45 | 305 |
|  | Number of patches of high vegetation | 1 | 2 | 14 | 93 |
|  | Shannon diversity index | 0.61 | 1.12 | 1.35 | 1.51 |
| BG02 | % low vegetation | 13.88 | 20.34 | 22.10 | 20.99 |
|  | % high vegetation | 86.12 | 70.95 | 55.66 | 38.64 |
|  | % buildings | 0 | 1.31 | 8.29 | 13.59 |
|  | % roads | 0 | 1.55 | 4.60 | 9.28 |
|  | % others | 0 | 5.67 | 9.35 | 17.50 |
|  | Total edge length of low vegetation (m) | 66.5 | 817.5 | 3531.5 | 27063 |
|  | Total edge length of high vegetation (m) | 66.5 | 679 | 2386 | 17934.5 |
|  | Total edge length of buildings (m) | 0 | 54 | 958.5 | 9146 |
|  | Total edge of roads (m) | 0 | 69.5 | 684.5 | 8230 |
|  | Average size of a patch of low vegetation (Ha) | 0.009 | 0.011 | 0.021 | 0.015 |
|  | Average size of a patch of high vegetation (Ha) | 0.111 | 0.281 | 0.293 | 0.072 |
|  | Average size of a patch of buildings (Ha) | 0 | 0.006 | 0.017 | 0.027 |
|  | Average size of a patch of roads (Ha) | 0 | 0.001 | 0.049 | 0.107 |
|  | Number of patches of low vegetation | 2 | 14 | 33 | 278 |
|  | Number of patches of high vegetation | 1 | 2 | 6 | 105 |
|  | Shannon diversity index | 0.41 | 0.87 | 1.23 | 1.49 |
| BG03 | % low vegetation | 32.95 | 23.86 | 25.98 | 16.70 |
|  | % high vegetation | 66.14 | 75.17 | 50.94 | 25.82 |
|  | % buildings | 0 | 0 | 9.03 | 25.34 |
|  | % roads | 0 | 0 | 3.65 | 11.06 |
|  | % others | 0.82 | 0.97 | 10.40 | 21.08 |
|  | Total edge length of low vegetation (m) | 161 | 532.5 | 3571 | 19158.5 |
|  | Total edge length of high vegetation (m) | 147 | 462.5 | 2611 | 12742.5 |
|  | Total edge length of buildings (m) | 0 | 0 | 932 | 15161 |
|  | Total edge of roads (m) | 0 | 0 | 405.5 | 9353 |
|  | Average size of a patch of low vegetation (Ha) | 0.014 | 0.038 | 0.030 | 0.016 |
|  | Average size of a patch of high vegetation (Ha) | 0.085 | 0.597 | 0.052 | 0.038 |
|  | Average size of a patch of buildings (Ha) | 0 | 0 | 0.028 | 0.051 |
|  | Average size of a patch of roads (Ha) | 0 | 0 | 0.116 | 0.091 |
|  | Number of patches of low vegetation | 3 | 5 | 27 | 200 |
|  | Number of patches of high vegetation | 1 | 1 | 31 | 135 |
|  | Shannon diversity index | 0.68 | 0.60 | 1.27 | 1.57 |
| BG04 | % low vegetation | 19.53 | 35.11 | 25.50 | 17.65 |
|  | % high vegetation | 80.47 | 55.66 | 50.00 | 26.15 |
|  | % buildings | 0 | 3.79 | 8.96 | 24.02 |
|  | % roads | 0 | 0 | 0.75 | 10.12 |
|  | % others | 0 | 5.44 | 14.79 | 22.06 |
|  | Total edge length of low vegetation (m) | 126.5 | 948 | 3400.5 | 19870 |
|  | Total edge length of high vegetation (m) | 126.5 | 848.5 | 2508.5 | 12823 |
|  | Total edge length of buildings (m) | 0 | 97.5 | 1126.5 | 14276.5 |
|  | Total edge of roads (m) | 0 | 0 | 94.5 | 8483 |
|  | Average size of a patch of low vegetation (Ha) | 0.005 | 0.034 | 0.025 | 0.018 |
|  | Average size of a patch of high vegetation (Ha) | 0.103 | 0.073 | 0.105 | 0.037 |
|  | Average size of a patch of buildings (Ha) | 0 | 0.015 | 0.029 | 0.054 |
|  | Average size of a patch of roads (Ha) | 0 | 0 | 0.026 | 0.071 |
|  | Number of patches of low vegetation | 5 | 8 | 32 | 189 |
|  | Number of patches of high vegetation | 1 | 6 | 15 | 139 |
|  | Shannon diversity index | 0.50 | 0.98 | 1.23 | 1.56 |
| BG05 | % low vegetation | 12.63 | 31.10 | 19.68 | 14.84 |
|  | % high vegetation | 87.37 | 50.43 | 29.59 | 26.11 |
|  | % buildings | 0 | 0 | 19.22 | 25.57 |
|  | % roads | 0 | 9.42 | 10.81 | 11.95 |
|  | % others | 0 | 9.05 | 20.70 | 21.53 |
|  | Total edge length of low vegetation (m) | 100.5 | 1268.5 | 3543.5 | 16883.5 |
|  | Total edge length of high vegetation (m) | 100.5 | 1064.5 | 2895 | 11488 |
|  | Total edge length of buildings (m) | 0 | 0 | 1886.5 | 13400 |
|  | Total edge of roads (m) | 0 | 320 | 1592 | 9053 |
|  | Average size of a patch of low vegetation (Ha) | 0.008 | 0.022 | 0.013 | 0.017 |
|  | Average size of a patch of high vegetation (Ha) | 0.038 | 0.031 | 0.024 | 0.044 |
|  | Average size of a patch of buildings (Ha) | 0 | 0 | 0.028 | 0.052 |
|  | Average size of a patch of roads (Ha) | 0 | 0.038 | 0.068 | 0.124 |
|  | Number of patches of low vegetation | 2 | 11 | 47 | 170 |
|  | Number of patches of high vegetation | 3 | 13 | 39 | 118 |
|  | Shannon diversity index | 0.38 | 1.15 | 1.56 | 1.57 |
| BG11 | % low vegetation | 39.07 | 21.76 | 21.14 | 17.22 |
|  | % high vegetation | 0 | 5.5 | 13.19 | 18.98 |
|  | % buildings | 23.52 | 21.15 | 23.36 | 16.00 |
|  | % roads | 23.48 | 27.24 | 17.57 | 13.00 |
|  | % others | 13.93 | 24.35 | 24.74 | 34.80 |
|  | Total edge length of low vegetation (m) | 215.5 | 1023 | 3690.5 | 21726 |
|  | Total edge length of high vegetation (m) | 0 | 191 | 1381.5 | 11449.5 |
|  | Total edge length of buildings (m) | 99 | 627 | 2352.5 | 10267.5 |
|  | Total edge of roads (m) | 133 | 657.5 | 1621 | 10163 |
|  | Average size of a patch of low vegetation (Ha) | 0.025 | 0.017 | 0.016 | 0.015 |
|  | Average size of a patch of high vegetation (Ha) | 0 | 0.008 | 0.028 | 0.032 |
|  | Average size of a patch of buildings (Ha) | 0.015 | 0.024 | 0.041 | 0.040 |
|  | Average size of a patch of roads (Ha) | 0.030 | 0.216 | 0.139 | 0.096 |
|  | Number of patches of low vegetation | 2 | 10 | 42 | 223 |
|  | Number of patches of high vegetation | 0 | 6 | 15 | 115 |
|  | Shannon diversity index | 1.32 | 1.52 | 1.59 | 1.54 |
| BG12_13 | % low vegetation | 39.35 | 37.92 | 36.47 | 27.30 |
|  | % high vegetation | 3.22 | 9.88 | 10.15 | 23.85 |
|  | % buildings | 16.32 | 16.61 | 17.38 | 16.46 |
|  | % roads | 26.42 | 14.28 | 15.34 | 10.22 |
|  | % others | 14.69 | 21.31 | 20.66 | 22.17 |
|  | Total edge length of low vegetation (m) | 240.5 | 1286 | 4356 | 30965 |
|  | Total edge length of high vegetation (m) | 40 | 385 | 917 | 14012 |
|  | Total edge length of buildings (m) | 128 | 705 | 2699 | 14626 |
|  | Total edge of roads (m) | 153 | 570 | 2240.5 | 10087.5 |
|  | Average size of a patch of low vegetation (Ha) | 0.012 | 0.027 | 0.024 | 0.020 |
|  | Average size of a patch of high vegetation (Ha) | 0.001 | 0.011 | 0.015 | 0.050 |
|  | Average size of a patch of buildings (Ha) | 0.005 | 0.010 | 0.015 | 0.019 |
|  | Average size of a patch of roads (Ha) | 0.034 | 0.039 | 0.081 | 0.067 |
|  | Number of patches of low vegetation | 4 | 11 | 48 | 272 |
|  | Number of patches of high vegetation | 3 | 8 | 21 | 93 |
|  | Shannon diversity index | 1.41 | 1.52 | 1.52 | 1.56 |
| BG14 | % low vegetation | 18.55 | 26.90 | 23.23 | 17.00 |
|  | % high vegetation | 59.86 | 35.56 | 15.66 | 17.26 |
|  | % buildings | 16.58 | 18.63 | 21.07 | 19.34 |
|  | % roads | 0 | 7.02 | 12.58 | 15.82 |
|  | % others | 5.01 | 11.89 | 27.46 | 30.58 |
|  | Total edge length of low vegetation (m) | 149.5 | 1270 | 3786.5 | 21961 |
|  | Total edge length of high vegetation (m) | 102.5 | 683.5 | 1525 | 11540.5 |
|  | Total edge length of buildings (m) | 57 | 512.5 | 2438.5 | 12345.5 |
|  | Total edge of roads (m) | 0 | 278.5 | 1827 | 12118.5 |
|  | Average size of a patch of low vegetation (Ha) | 0.008 | 0.014 | 0.028 | 0.015 |
|  | Average size of a patch of high vegetation (Ha) | 0.019 | 0.056 | 0.023 | 0.026 |
|  | Average size of a patch of buildings (Ha) | 0.022 | 0.037 | 0.039 | 0.044 |
|  | Average size of a patch of roads (Ha) | 0 | 0.028 | 0.031 | 0.130 |
|  | Number of patches of low vegetation | 3 | 15 | 26 | 228 |
|  | Number of patches of high vegetation | 4 | 5 | 21 | 130 |
|  | Shannon diversity index | 1.07 | 1.48 | 1.57 | 1.57 |
| BG15_16 | % low vegetation | 16.00 | 31.04 | 27.10 | 26.14 |
|  | % high vegetation | 11.32 | 17.51 | 12.67 | 15.52 |
|  | % buildings | 23.62 | 20.89 | 20.03 | 17.68 |
|  | % roads | 30.12 | 13.16 | 19.31 | 14.98 |
|  | % others | 18.94 | 16.60 | 20.89 | 25.68 |
|  | Total edge length of low vegetation (m) | 169.5 | 1218 | 4283.5 | 24617.5 |
|  | Total edge length of high vegetation (m) | 70.5 | 507 | 1553.5 | 11049 |
|  | Total edge length of buildings (m) | 135 | 806 | 2723.5 | 14763 |
|  | Total edge of roads (m) | 156.5 | 498.5 | 2333.5 | 12141.5 |
|  | Average size of a patch of low vegetation (Ha) | 0.004 | 0.022 | 0.018 | 0.023 |
|  | Average size of a patch of high vegetation (Ha) | 0.012 | 0.023 | 0.016 | 0.026 |
|  | Average size of a patch of buildings (Ha) | 0.010 | 0.014 | 0.017 | 0.020 |
|  | Average size of a patch of roads (Ha) | 0.033 | 0.032 | 0.075 | 0.109 |
|  | Number of patches of low vegetation | 5 | 12 | 48 | 221 |
|  | Number of patches of high vegetation | 2 | 6 | 25 | 116 |
|  | Shannon diversity index | 1.59 | 1.55 | 1.58 | 1.58 |
| BG21 | % low vegetation | 38.01 | 12.40 | 15.00 | 13.37 |
|  | % high vegetation | 30.01 | 24.16 | 24.88 | 18.86 |
|  | % buildings | 0 | 2.50 | 20.31 | 28.80 |
|  | % roads | 1.50 | 15.26 | 10.33 | 12.95 |
|  | % others | 30.48 | 45.68 | 29.48 | 26.02 |
|  | Total edge length of low vegetation (m) | 253 | 738 | 3379 | 17504 |
|  | Total edge length of high vegetation (m) | 162.5 | 425.5 | 2059.5 | 10541.5 |
|  | Total edge length of buildings (m) | 0 | 68 | 1703.5 | 16761 |
|  | Total edge of roads (m) | 18.5 | 356 | 1185 | 9767.5 |
|  | Average size of a patch of low vegetation (Ha) | 0.025 | 0.012 | 0.014 | 0.015 |
|  | Average size of a patch of high vegetation (Ha) | 0.008 | 0.027 | 0.031 | 0.030 |
|  | Average size of a patch of buildings (Ha) | 0 | 0.020 | 0.049 | 0.056 |
|  | Average size of a patch of roads (Ha) | 0.002 | 0.122 | 0.046 | 0.098 |
|  | Number of patches of low vegetation | 2 | 8 | 33 | 172 |
|  | Number of patches of high vegetation | 5 | 7 | 26 | 123 |
|  | Shannon diversity index | 1.16 | 1.34 | 1.55 | 1.55 |
| BG22 | % low vegetation | 9.65 | 9.41 | 6.64 | 9.94 |
|  | % high vegetation | 28.40 | 23.53 | 14.2 | 17.97 |
|  | % buildings | 29.65 | 33.66 | 44.99 | 37.18 |
|  | % roads | 0 | 10.86 | 12.40 | 13.52 |
|  | % others | 32.45 | 22.50 | 21.79 | 21.39 |
|  | Total edge length of low vegetation (m) | 186.5 | 807 | 1980.5 | 12947 |
|  | Total edge length of high vegetation (m) | 109.5 | 500 | 1397 | 8694.5 |
|  | Total edge length of buildings (m) | 95.5 | 752.5 | 3400 | 18334 |
|  | Total edge of roads (m) | 1 | 353 | 1990.5 | 10921.5 |
|  | Average size of a patch of low vegetation (Ha) | 0.002 | 0.004 | 0.006 | 0.012 |
|  | Average size of a patch of high vegetation (Ha) | 0.018 | 0.031 | 0.025 | 0.033 |
|  | Average size of a patch of buildings (Ha) | 0.018 | 0.024 | 0.059 | 0.072 |
|  | Average size of a patch of roads (Ha) | 0 | 0.086 | 0.056 | 0.178 |
|  | Number of patches of low vegetation | 5 | 18 | 36 | 161 |
|  | Number of patches of high vegetation | 2 | 6 | 18 | 107 |
|  | Shannon diversity index | 1.31 | 1.51 | 1.41 | 1.51 |
| BG23 | % low vegetation | 0 | 2.50 | 7.50 | 13.14 |
|  | % high vegetation | 2.67 | 5.47 | 10.19 | 21.69 |
|  | % buildings | 41.74 | 46.34 | 39.69 | 23.28 |
|  | % roads | 18.34 | 21.30 | 20.26 | 13.69 |
|  | % others | 37.29 | 24.39 | 22.36 | 28.20 |
|  | Total edge length of low vegetation (m) | 0 | 232 | 2027 | 18469.5 |
|  | Total edge length of high vegetation (m) | 18.5 | 164 | 1232 | 11018 |
|  | Total edge length of buildings (m) | 160 | 1150 | 4092 | 14429 |
|  | Total edge of roads (m) | 108 | 766 | 2652 | 10268.5 |
|  | Average size of a patch of low vegetation (Ha) | 0 | 0.001 | 0.006 | 0.011 |
|  | Average size of a patch of high vegetation (Ha) | 0.003 | 0.022 | 0.016 | 0.033 |
|  | Average size of a patch of buildings (Ha) | 0.018 | 0.023 | 0.044 | 0.048 |
|  | Average size of a patch of roads (Ha) | 0.024 | 0.169 | 0.058 | 0.149 |
|  | Number of patches of low vegetation | 0 | 15 | 43 | 228 |
|  | Number of patches of high vegetation | 1 | 2 | 20 | 128 |
|  | Shannon diversity index | 1.14 | 1.28 | 1.45 | 1.57 |
| BG24 | % low vegetation | 25.49 | 15.87 | 14.73 | 19 |
|  | % high vegetation | 17.88 | 16.74 | 17.17 | 16.3 |
|  | % buildings | 21.31 | 23.94 | 19.70 | 20.6 |
|  | % roads | 0 | 18.69 | 16 | 13.3 |
|  | % others | 35.31 | 24.76 | 32.4 | 30.7 |
|  | Total edge length of low vegetation (m) | 287.5 | 1214.5 | 3471.5 | 19526 |
|  | Total edge length of high vegetation (m) | 138 | 698.5 | 2195.5 | 10587 |
|  | Total edge length of buildings (m) | 113 | 734 | 2141.5 | 12711.5 |
|  | Total edge of roads (m) | 0 | 514.5 | 1795.5 | 8381.5 |
|  | Average size of a patch of low vegetation (Ha) | 0.016 | 0.009 | 0.012 | 0.017 |
|  | Average size of a patch of high vegetation (Ha) | 0.006 | 0.007 | 0.013 | 0.022 |
|  | Average size of a patch of buildings (Ha) | 0.007 | 0.017 | 0.025 | 0.035 |
|  | Average size of a patch of roads (Ha) | 0 | 0.074 | 0.046 | 0.119 |
|  | Number of patches of low vegetation | 2 | 15 | 40 | 213 |
|  | Number of patches of high vegetation | 4 | 20 | 40 | 146 |
|  | Shannon diversity index | 1.35 | 1.59 | 1.56 | 1.57 |
