## Supplementary material for "Investigating the role of urban vegetation alongside other environmental variables in shaping *Aedes albopictus* presence and abundance in Montpellier, France": S4_Table

| **Area** | **Mosquito female/trap/24h (± SE)** | **Mosquito male/trap/24h**  **(± SE)** | ***Ae. albopictus* female/trap/24h**  **(± SE)** | ***Ae. albopictus* male /trap/24h**  **(± SE)** | ***Cx. pipiens* female /trap/24h**  **(± SE)** | ***Cx. pipiens* male /trap/24h**  **(± SE)** | ***Cu. longiareolata* female /trap/24h**  **(± SE)** | ***Cu. longiareolata* male /trap/24h**  **(± SE)** | ***Cu. annulata* male /trap/24h**  **(± SE)** |
| --- | --- | --- | --- | --- | --- | --- | --- | --- | --- |
| PRK-AGL | 8.40 (± 3.18) | 2.58 (± 1.32) | 6.22 (± 3.05) | 1.64 (± 1.39) | 1.07 (± 0.43) | 0.11 (± 0.08) | 0.02 (± 0.02) | 0.02 (± 0.02) | 0 |
| PRK-BOT | 5.61 (± 1.96) | 5.15 (± 2.50) | 2.91 (± 1.36) | 2.10 (± 1.72) | 2.03 (± 0.69) | 1.68 (± 0.58) | 0.02 (± 0.02) | 0.02 (± 0.02) | 0.02 (± 0.02) |
| RES-AGR | 3.90 (± 1.51) | 1.82 (± 0.94) | 3.30 (± 1.64) | 1.18 (± 1.00) | 0.43 (± 0.19) | 0.04 (± 0.04) | 0 | 0 | 0 |
| RES-LEM | 3.83 (± 1.51) | 1.81 (± 0.97) | 3.31 (± 1.68) | 0.92 (± 0.79) | 0.39 (± 0.18) | 0.26 (± 0.15) | 0 | 0.02 (± 0.02) | 0 |
| RES-SOUL | 4.17 (± 1.60) | 1.39 (± 0.73) | 3.29 (± 1.63) | 0.88 (± 0.74) | 0.25 (± 0.13) | 0.04 (± 0.04) | 0.02 (± 0.02) | 0 | 0 |
| IMP-ACA | 5.84 (± 2.67) | 5.35 (± 3.08) | 4.71 (± 2.65) | 3.34 (± 3.04) | 0.48 (± 0.28) | 0.30 (± 0.22) | 0 | 0 | 0 |
| IMP-BIB | 1.08 (± 0.53) | 0.52 (± 0.34) | 0.92 (± 0.57) | 0.26 (±0.2) | 0 | 0.08 (± 0.08) | 0 | 0.03 (± 0.03) | 0 |
| IMP-DID | 0.61 (± 0.34) | 0.04 (± 0.04) | 0.45 (± 0.30) | 0.03 (± 0.03) | 0 | 0 | 0.04 (± 0.04) | 0 | 0 |
| IMP-SCU | 13.7 (± 6.24) | 6.17 (± 3.54) | 5.56 (± 3.14) | 3.94 (± 3.58) | 2.56 (± 1.19) | 0.69(± 0.43) | 0 | 0 | 0 |

The mosquito density represents the marginal mean of the number of mosquitoes caught per trap per 24 hours, calculated based on the sampling environment, with the standard error specified (SE).
