## Supplementary material for "Investigating the role of urban vegetation alongside other environmental variables in shaping *Aedes albopictus* presence and abundance in Montpellier, France": S5_Table

| **Analysis** | **Variable topic** | **Variable code** | **Variable name** | **Selected for multivariate analysis (Yes/No)** | **Reason for exclusion** |
| --- | --- | --- | --- | --- | --- |
| Presence | Weeks-lagged meteorological | TMN_1_1 | Daily average temperature during the first week before sampling | No | Correlated with TMIN_1_1, TMAX_1_1, tmean_collection, GDD_1_1, RH_6_6 |
| Presence | Weeks-lagged meteorological | TMIN_1_1 | Daily minimum temperature during the first week before sampling | No | Correlated with TMN_1_1, TMAX_0_1, RH_6_6, GDD_1_1, tmean_collection |
| Presence | Weeks-lagged meteorological | TMAX_0_1 | Daily maximum temperature between the sampling and the first week before | No | Correlated with TMN_1_1, TMIN_1_1, RH_6_6, GDD_1_1, tmean_collection |
| Presence | Weeks-lagged meteorological | GDD_1_1 | Weekly cumulated Growing Deegree Day during first week before sampling | No | Although this variable is correlated with others (TMN_1_1, TMIN_1_1, RH_6_6, TMAX_0_1), it is recognized as one of the key factors driving *Aedes albopictus* population dynamics (Roiz et al. 2010). It was initially retained during the preliminary selection process for correlated variables but was ultimately excluded due to a Variance Inflation Factor (VIF) exceeding 3. |
| Presence | Weeks-lagged meteorological | RH_6_6 | Daily relative humidity during the six week before sampling | No | Correlated with TMN_1_1, TMIN_1_1, TMAX_0_1, GDD_1_1, tmean_collection |
| Presence | Weeks-lagged meteorological | CUMRF_6_6 | Weekly cumulated rainfall during the six week before sampling | Yes | *NA* |
| Presence | Weeks-lagged air quality | PM2.5_0_1 | Daily concentration of PM2.5 during the sampling and the first week before | No | Correlated with PM10_0_1 and with all weeks-lagged temperature. |
| Presence | Weeks-lagged air quality | PM10_0_1 | Daily concentration of PM10 during the sampling and the first week before | No | Correlated with PM2.5_0_1 and with all weeks-lagged temperature. |
| Presence | Weeks-lagged air quality | O3_3_3 | Daily concentration of O3 during the third week before sampling | No | Correlated with weeks-lagged of rainfall and wind |
| Presence | Weeks-lagged air quality | NO2_0_0 | Daily concentration of NO2 during the sampling | Yes | *NA* |
| Presence | Real-time and 48h-lagged microclimatic | rhmean_24h_48h | Average hourly relative humidity between 24h and 48h before sampling | Yes | *NA* |
| Presence | Real-time and 48h-lagged microclimatic | rhmax_24h_48h | Maximum hourly relative humidity between 24h and 48h before sampling | No | Correlated with rhmean_24h_48h and VIF>3 |
| Presence | Real-time and 48h-lagged microclimatic | rhmin_collection | Minimum hourly relative humidity during 24h of sampling | No | Correlated with Rainfall_Collection and VIF>3 |
| Presence | Real-time and 48h-lagged microclimatic | tmean_collection | Average hourly temperature during the 24h of sampling | No | Correlated with TMN_1_1, TMIN_1_1, TMAX_0_1, GDD_1_1, tmin_24h_48h, and tmax_collection. |
| Presence | Real-time and 48h-lagged microclimatic | tmax_collection | Maximum hourly temperature during the 24h of sampling | No | Correlated with TMAX_0_1, tmin_24h_48h, and tmean_collection and VIF>3. |
| Presence | Real-time and 48h-lagged microclimatic | tmin_24h_48h | Minimum hourly temperature between 24h and 48h before sampling | Yes | *NA* |
| Presence | Real-time and 48h-lagged meteorological | Rainfall_collection | Rainfall cumulated during the 24h of sampling | No | Correlated with rhmin_collection and VIF>3 |
| Presence | Real-time and 48h-lagged meteorological | Patmin_collection | Minimum atmospheric pressure during the 24h of sampling | Yes | *NA* |
| Presence | Real-time and 48h-lagged meteorological | Patmean_collection | Average atmospheric pressure during the 24h of sampling | No | Correlated with Patmin_collection and Patmax_collection and VIF>3. |
| Presence | Real-time and 48h-lagged meteorological | Patmax_collection | Maximum atmospheric pressure during the 24h of sampling | No | Correlated with Patmin_collection and Patmean_collection and VIF>3. |
| Presence | Real-time and 48h-lagged meteorological | Patdiff_collection | Difference of mean daily atmospheric pressure between the sampling day and the previous day | Yes | *NA* |
| Presence | Land cover | Edge_high_veget_20 | Total length of edge of high vegetation at 20 m | No | Correlated with Edge_roads_20 and Area_roads_20 and VIF>3. |
| Presence | Land cover | Edge_roads_20 | Total length of edge of roads at 20 m | Yes | *NA* |
| Presence | Land cover | Area_roads_20 | Average patch size of roads at 20 m | No | Correlated with Edge_buildings_20, Edge_roads_20 and Edge_high_veget_20, and VIF>3. |
| Presence | Land cover | Edge_buildings_20 | Total length of edge of buildings at 20 m | No | Correlated with Edge_roads_20, Area_roads_20 and %_buildings_50, and VIF>3. |
| Presence | Land cover | %_buildings_50 | Percentage of cover of buildings at 50 m | No | Correlated with Edge_buildings_20 and VIF>3. |
| Presence | Land cover | Area_low_veget_20 | Average patch size of low vegetation at 20 m | Yes | *NA* |
| Presence | Land cover | Shannon index | Shannon diversity indes | No | VIF>3 |
| Presence | Land cover | Patches_high_veget_20 | Number of patches of high vegetation at 20 m | Yes | *NA* |
| Presence | Land cover | Edge_low_veget_20 | lsm_c_pland_LCG_20_13, | Yes | *NA* |
| Presence | Land cover | %_low_veget_20 | lsm_c_te_LCG_20_13, lsm_c_area_mn_LCG_20_13 | No | Correlated with Area_low_veget_20 and Edge_low_veget_20 and VIF>3. |
| Presence | Demographic | POP_250 | Population density at 250 m | No | Correlated with Pov_Hous and VIF>3. |
| Presence | Socio-economic | Pov_Hous | Numer of poor households | No | Excluded during the recursive elimination features |
| Abundance | Weeks-lagged meteorological | TMN_0_2 | Daily average temperature during the sampling and the second week before | No | Correlated with GDD_0_2, TMIN_0_2, TMAX_0_2, rhmean_collection, tmean_collection, tmin_collection, RH_6_6 |
| Abundance | Weeks-lagged meteorological | TMIN_0_2 | Daily minimum temperature during the sampling and the second week before | No | Correlated with GDD_0_2, TMN_0_2, TMAX_0_2, rhmean_collection, tmean_collection, tmin_collection, RH_6_6 |
| Abundance | Weeks-lagged meteorological | TMAX_0_2 | Daily maximum temperature during the sampling and the second week before | No | Correlated with GDD_0_2, TMIN_0_2, TMN_0_2, rhmean_collection, tmean_collection, tmin_collection, RH_6_6 |
| Abundance | Weeks-lagged meteorological | GDD_0_2 | Weekly cumulated Growing Deegree Day during the sampling and the second week before | Yes | *NA* |
| Abundance | Weeks-lagged meteorological | RH_6_6 | Daily relative humidity during the six week before sampling | No | Correlated with GDD_0_2, TMIN_0_2, TMAX_0_2, rhmean_collection, tmean_collection, tmin_collection. |
| Abundance | Weeks-lagged meteorological | CUMRF_6_6 | Weekly cumulated rainfall during the six week before sampling | Yes | Excluded during the recursive elimination features but keeped for spatial autocorrelation |
| Abundance | Weeks-lagged meteorological | WS_0_5 | Mean daily wind speed between the sampling and the 5th week before | Yes | *NA* |
| Abundance | Weeks-lagged air quality | PM2.5_0_3 | Daily concentration of PM2.5 during the sampling and the third week before | No | Correlated with PM10_0_1 and with all weeks-lagged temperature. |
| Abundance | Weeks-lagged air quality | PM10_0_1 | Daily concentration of PM10 during the sampling and the first week before | No | Correlated with PM2.5_0_3 and with all weeks-lagged temperature. |
| Abundance | Weeks-lagged air quality | O3_1_1 | Daily concentration of O3 during the first week before sampling | No | Correlated with weeks-lagged of rainfall. |
| Abundance | Real-time and 48h-lagged microclimatic | rhmean_collection | Average hourly relative humidity during 24h of sampling | No | Correlated with GDD_0_2, TMIN_0_2, TMN_0_2, TMAX_0_2, rhmin_collection |
| Abundance | Real-time and 48h-lagged microclimatic | rhmin_collection | Minimum hourly relative humidity during 24h of sampling | No | Correlated with rhmin_collection and VIF>3. |
| Abundance | Real-time and 48h-lagged microclimatic | rhmax_24h | Maximum hourly relative humidity during the 24h before sampling | Yes | Excluded during the recursive elimination features bit keeped because of spatial autocorrelation |
| Abundance | Real-time and 48h-lagged microclimatic | tmean_collection | Average hourly relative humidity during 24h of sampling | No | Correlated with GDD_0_2, TMIN_0_2, TMN_0_2, TMAX_0_2, tmin_collection, tmax_collection. |
| Abundance | Real-time and 48h-lagged microclimatic | tmin_collection | Minimum hourly relative humidity during 24h of sampling | No | Correlated with GDD_0_2, TMIN_0_2, TMN_0_2, TMAX_0_2, tmean_collection, tmax_collection. |
| Abundance | Real-time and 48h-lagged microclimatic | tmax_collection | Maximum hourly relative humidity during 24h of sampling | Yes | *NA* |
| Abundance | Real-time and 48h-lagged meteorological | Rainfall_24h | Cumulated rainfall during the 24h before sampling | No | Excluded during the recursive elimination features |
| Abundance | Real-time and 48h-lagged meteorological | Patmean_24h_48h | Average atmospheric pressure between the 24h and the 48h before sampling | No | Correlated with Patmax_24h_48h and Patdiff_24h_48h and VIF>3. |
| Abundance | Real-time and 48h-lagged meteorological | Patmax_24h_48h | Maximum atmospheric pressure between the 24h and the 48h before sampling | No | Correlated with Patmean_24h_48h and Patdiff_24h_48h and VIF>3. |
| Abundance | Real-time and 48h-lagged meteorological | Patdiff_24h_48h | Difference of daily atmospheric pressure between the 24h and 48h before sampling. | No | Excluded during the recursive elimination features |
| Abundance | Land cover | Patches_high_veget_50 | Number of patches of high vegetation at 50 m | No | VIF>3 |
| Abundance | Land cover | %_High_veget_20 | Percentage of cover of High vegetation at 20 m | No | Correlated with Edge_High_Veget_50, Edge_Buildings_100,  Environment (Park) and VIF>3 |
| Abundance | Land cover | Edge_High_Veget_50 | Total length of edges of high vegetation at 50 m | No | Correlated with %_High_veget_20 and VIF>3. |
| Abundance | Land cover | Area_High_Veget_100 | The average patch size of high vegetation at 100 m | No | Excluded during the recursive elimination features |
| Abundance | Land cover | Te_Low_Veget_100 | Total length of edges of low vegetation at 100 m | No | Correlated with %_low_veget_250 and Area_Roads_250, and VIF>3. |
| Abundance | Land cover | %_low_veget_250 | Percentage cover of low vegetation at 250 m | Yes | *NA* |
| Abundance | Land cover | Area_Low_Veget_50 | The average patch size of low vegetation at 50 m | Yes | *NA* |
| Abundance | Land cover | Edge_Roads_250 | The total length of roads edge at 250 m. | Yes | *NA* |
| Abundance | Land cover | %_Roads_250 | Percentage of cover of roads at 250 m. | No | Correlated with Edge_Roads_250 and VIF>3. |
| Abundance | Land cover | Area_Roads_250 | The average patch size of roads at 250 m | Yes | *NA* |
| Abundance | Land cover | Edge_Buildings_100 | The total length of buildings edge at 100 m. | No | Correlated with %_High_Veget_20, %_Building_100, Environment (Park) and VIF>3. |
| Abundance | Land cover | %_Buildings_100 | Percentage cover of buildings at 100 m | No | Correlated with Edge_Buildings_100, Area_Low_Veget_50, Area_Roads_250n and Area_Buildings_100, and VIF>3. |
| Abundance | Land cover | Area_Buildings_100 | The average patch size of buildings at 100 m | No | Correlated with %_Low_Veget_250, with %_Buildings_100, Number of poor households and Hom_Be_45 and VIF>3. |
| Abundance | Land cover | Environment (Park) | Urban park environment vs residential areas and built impervous areas | No | Correlated with %_High_Veget_20, %_Roads_250 and Edge_Buildings_100, and VIF>3. |
| Abundance | Socio-economic | Pov_Hou | Number of poor households | No | Excluded during the recursive elimination features |
| Abundance | Socio-economic | Hom_Be_45 | Homes built before 1945 | No | Correlated with %_Low_Veget_250 and Area_Buildings_100 and VIF>3. |
