## Supplementary figures and images for "Investigating the role of urban vegetation alongside other environmental variables in shaping *Aedes albopictus* presence and abundance in Montpellier, France"

### S1_Fig

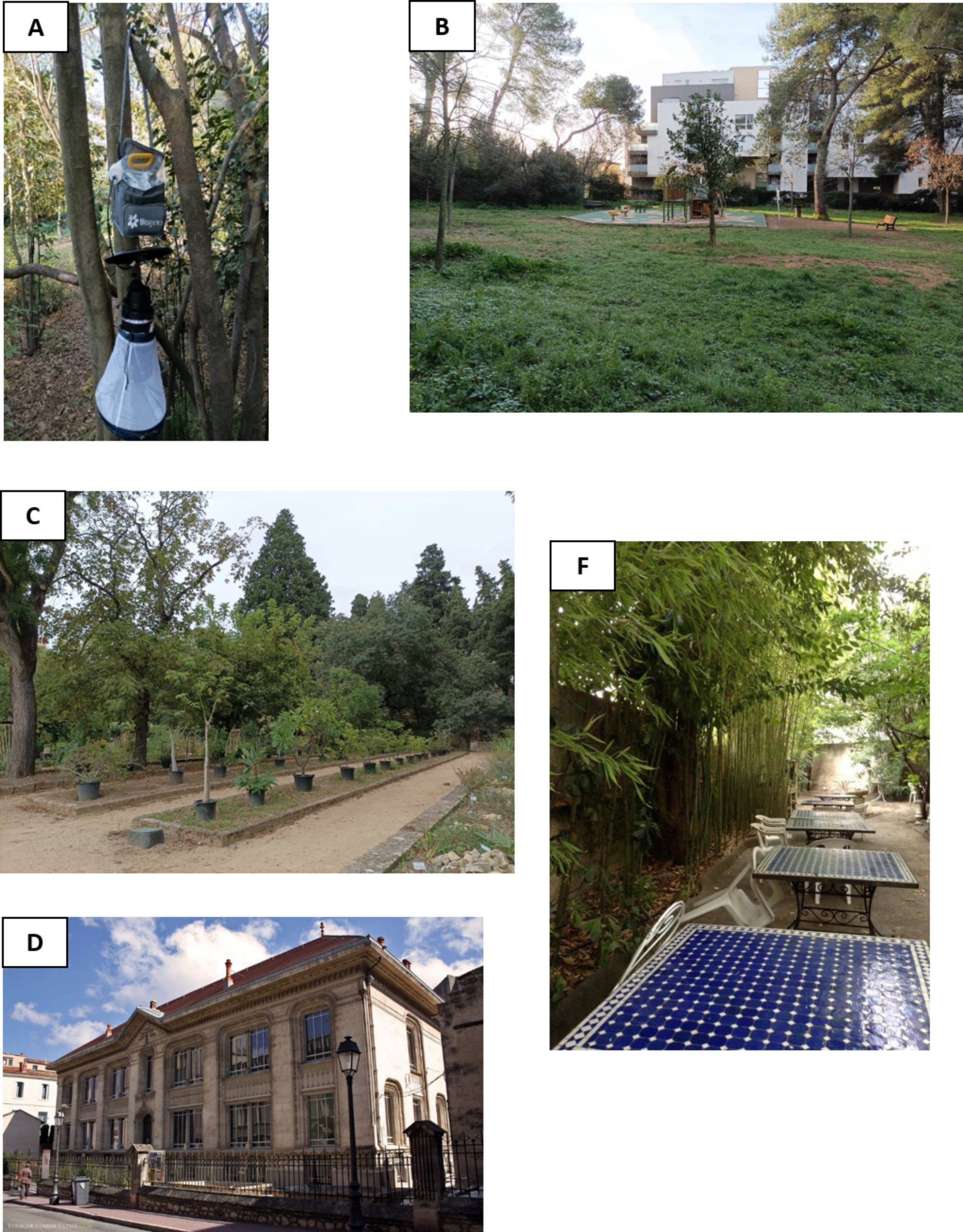

### S2_Fig

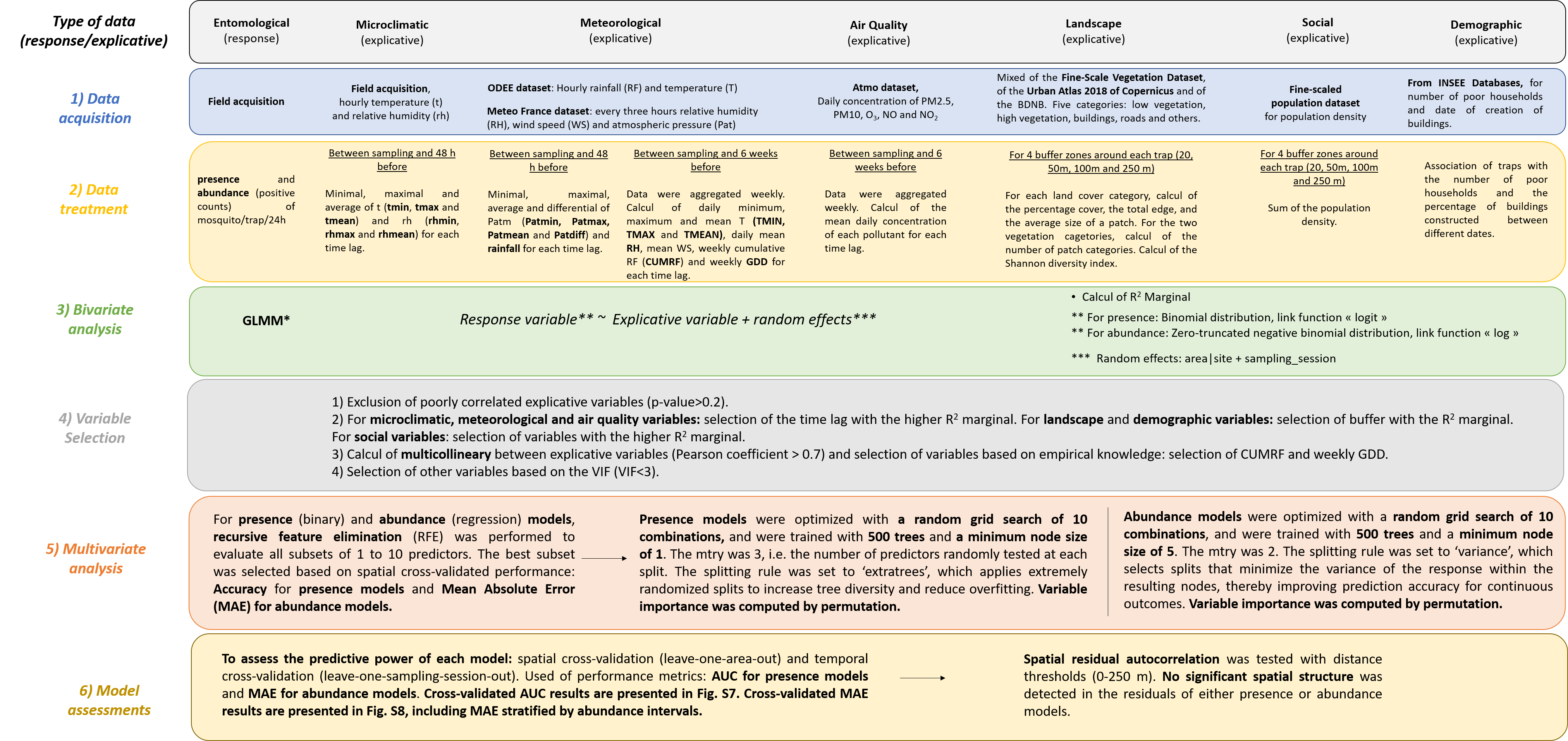

### S3_Fig

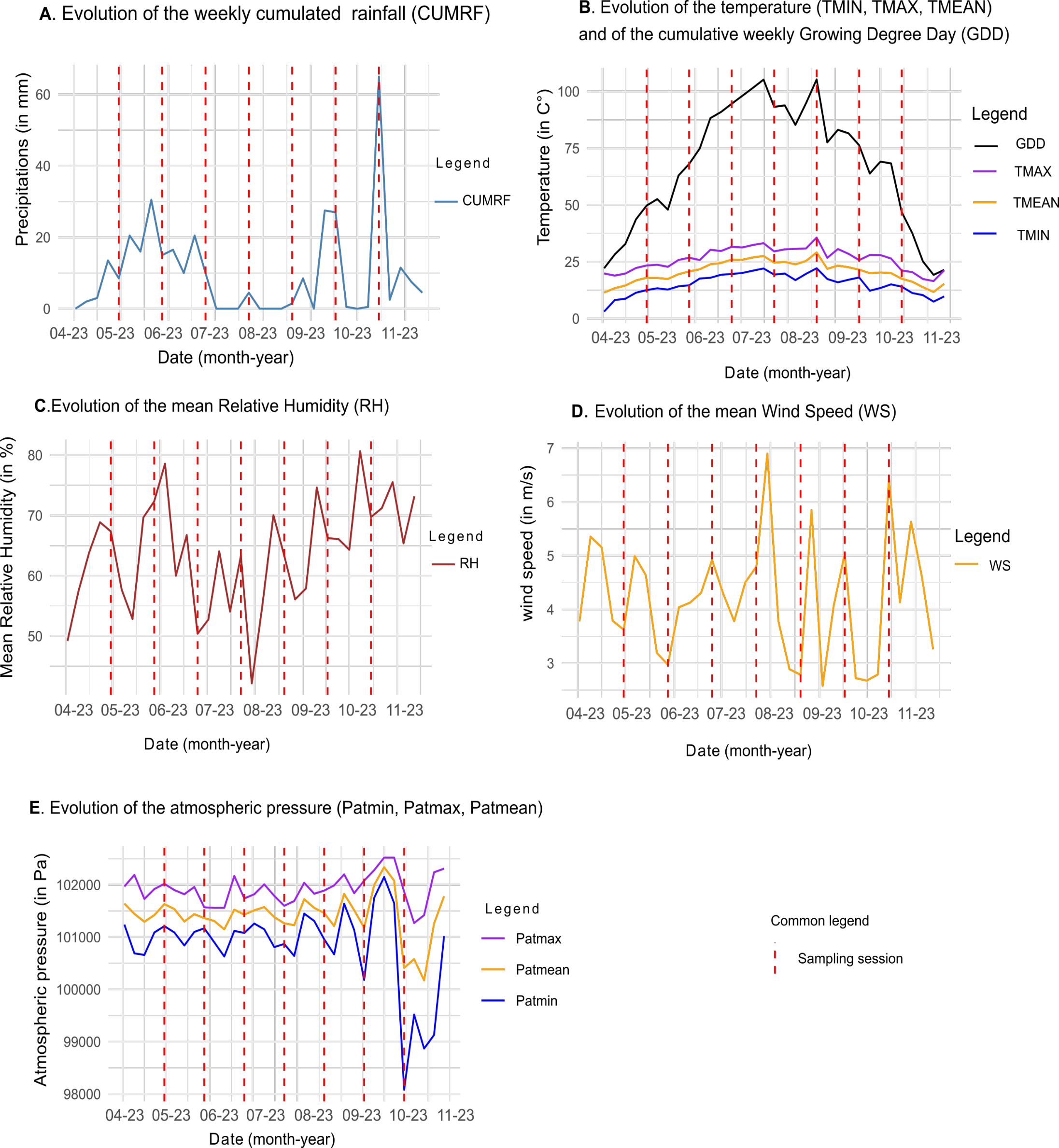

### S4_Fig

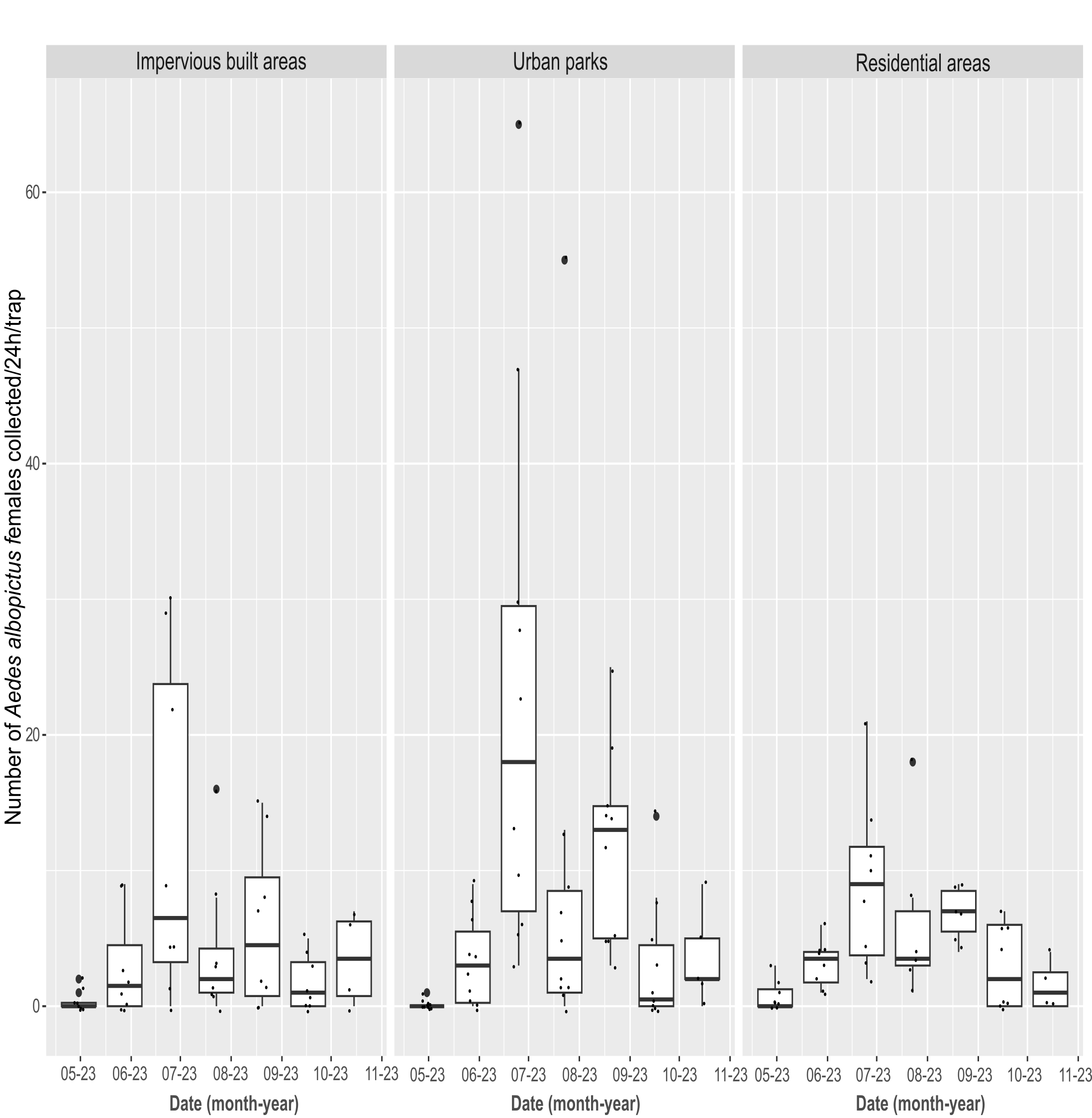

### S5_Fig

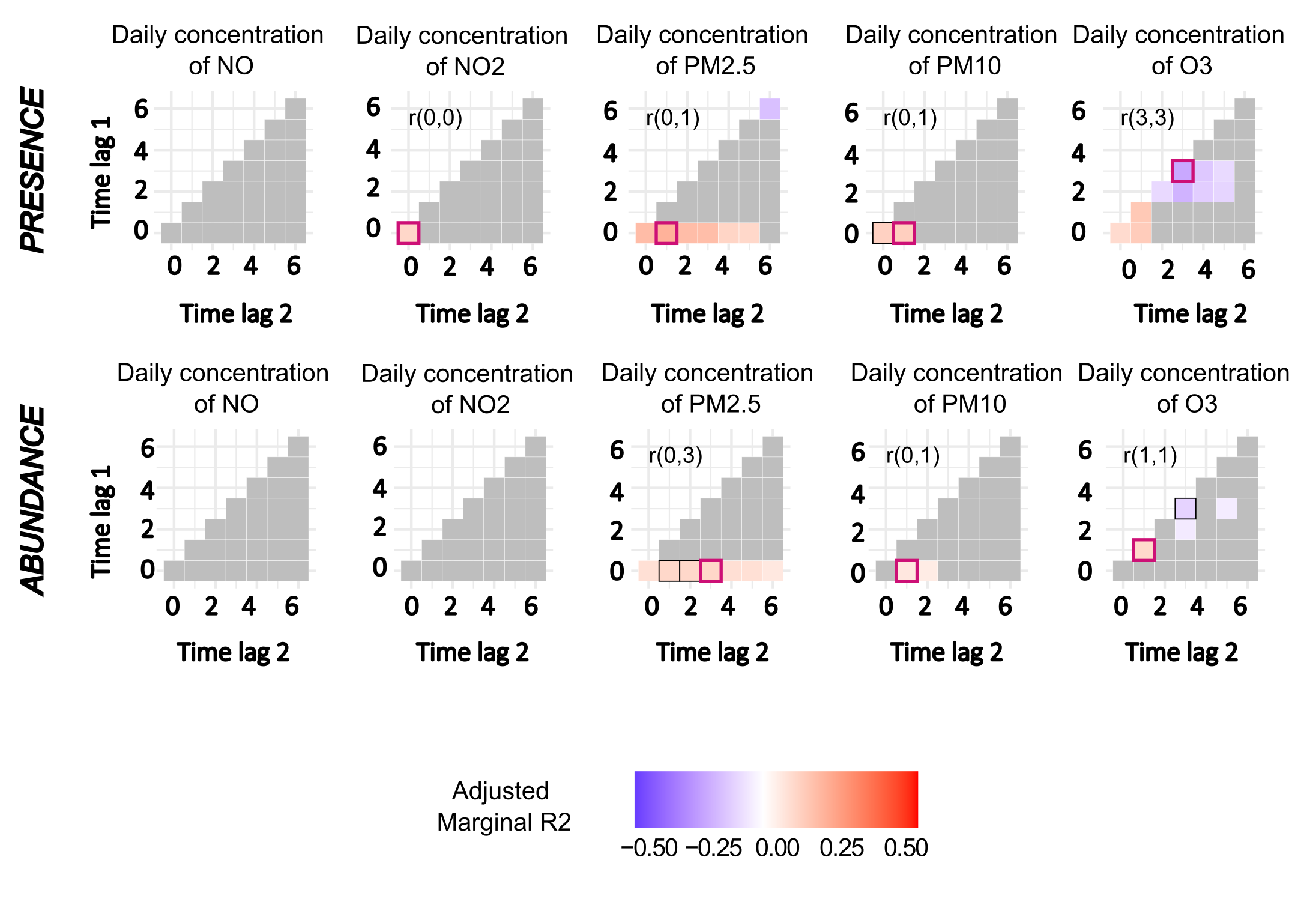

### S6_Fig

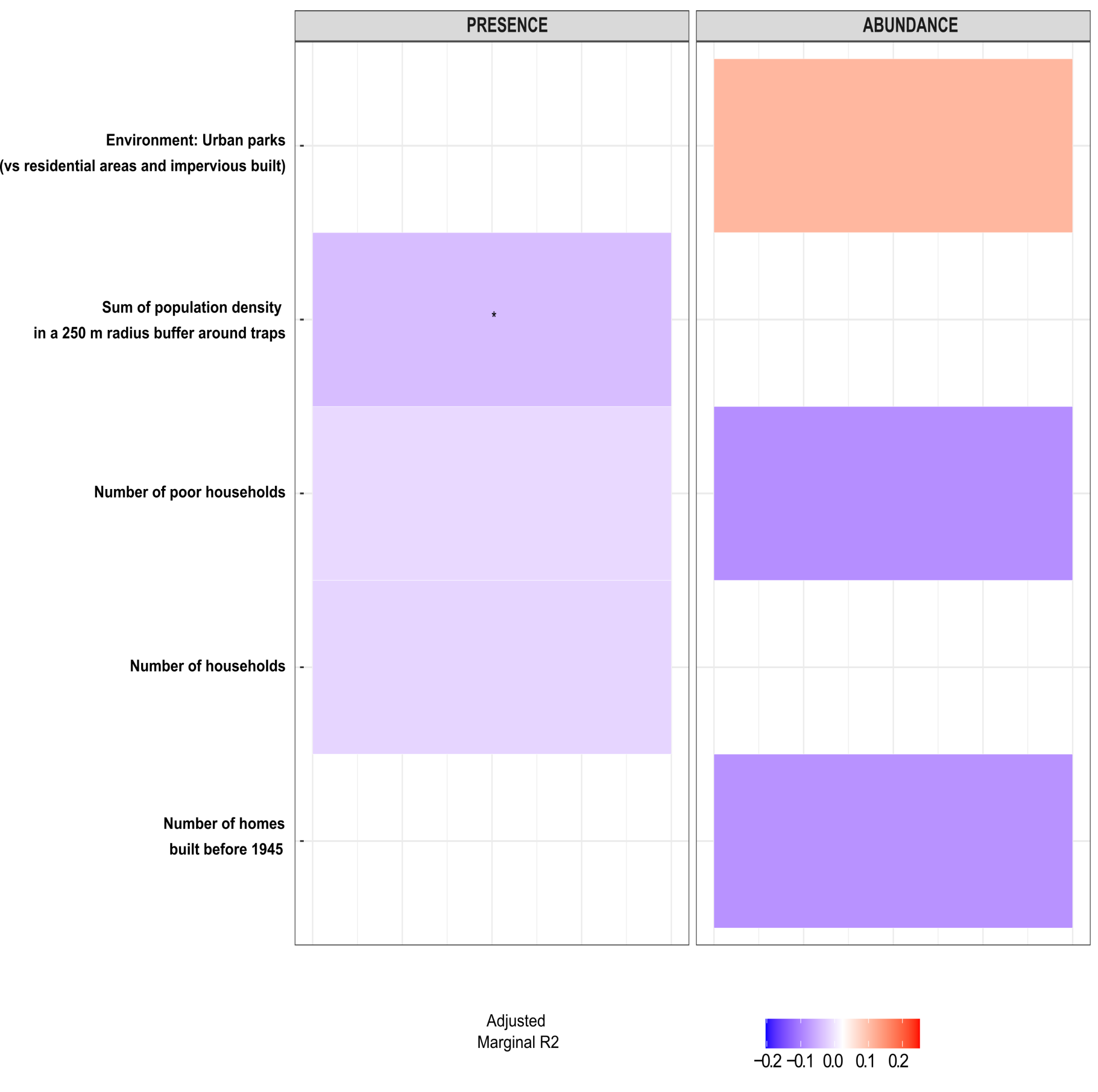

### S7_Fig

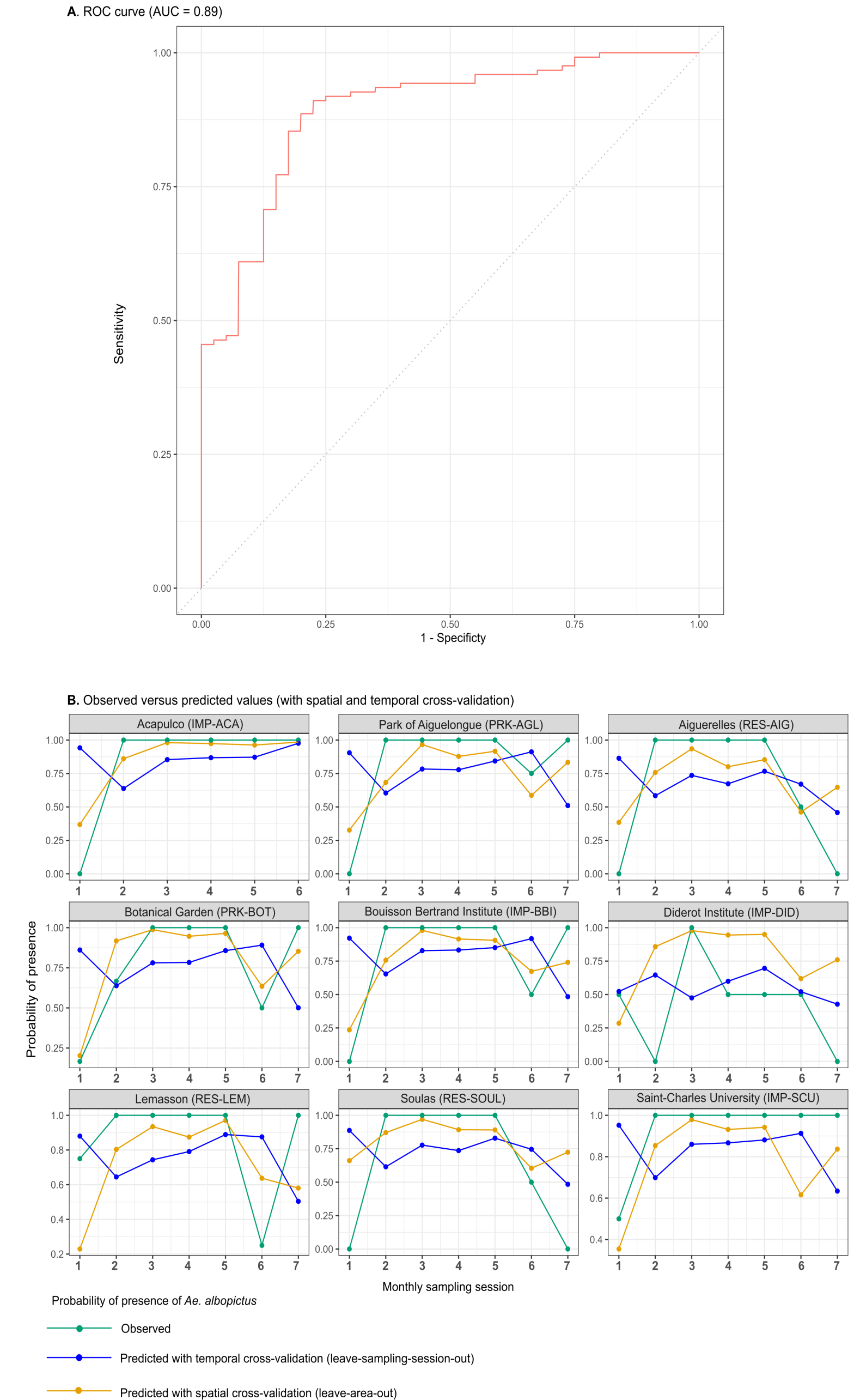

### S8_Fig

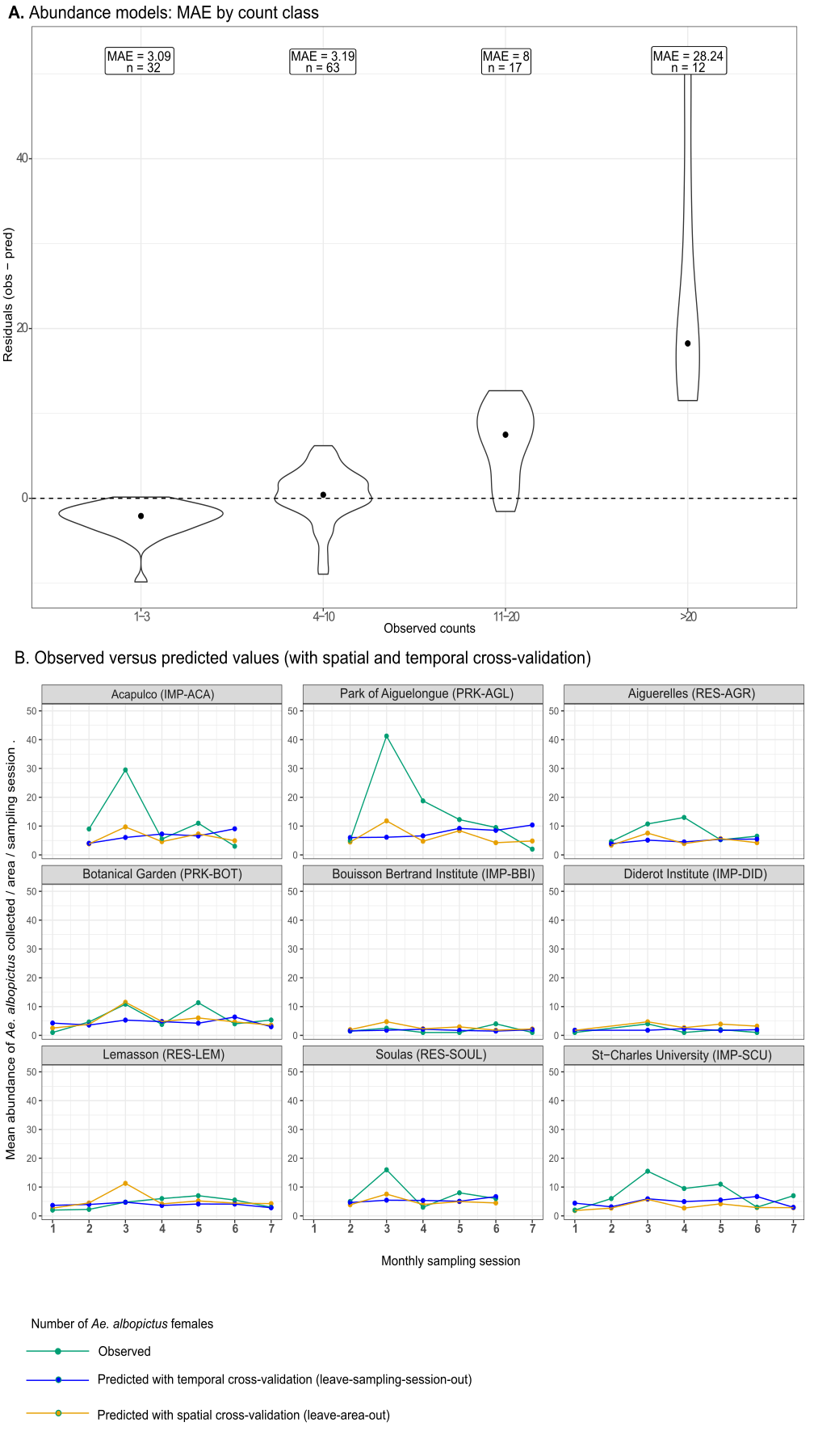
